# Ceg1 depletion reveals mechanisms governing degradation of non-capped RNAs

**DOI:** 10.1101/2022.09.24.509330

**Authors:** Onofrio Zanin, Daniel Hebenstreit, Pawel Grzechnik

**Affiliations:** School of Biosciences, University of Birmingham, Edgbaston, Birmingham, B15 2TT, UK; School of Life Sciences, University of Warwick, Coventry CV4 7AL, UK; School of Biological Sciences, University of Manchester, Oxford Road, Manchester M13 9PT, UK

**Keywords:** capping, RNA degradation, transcription

## Abstract

Most functional eukaryotic mRNAs contain a 7-methylguanosine (m7G) cap which serves as a platform that recruits proteins to support essential biological functions such as mRNA processing, nuclear export and translation. Although capping is accomplished during the first steps of transcription the fate and turnover of uncapped transcripts have not been studied extensively. Here, we employed fast nuclear depletion of the capping enzymes in *Saccharomyces cerevisiae* to uncover the turnover of the transcripts that failed to be capped. We show that the levels of non-capped mRNAs are determined principally by the abundance of their synthesis. Nuclear depletion of the capping enzymes increases the levels of lowly expressed mRNAs and decreases mRNAs that are highly transcribed altogether mimicking the effects observed in cells lacking the predominantly cytoplasmic 5’-3’ exonuclease Xrn1. The nuclear 5’-3’ exonuclease Rat1 is not involved in the degradation of cap-defective transcripts and the lack of the capping does not affect the distribution of RNA Polymerase II on the chromatin. Our data indicate that the mRNAs that failed to be capped are not directed to a specific quality-control pathway and that the 5’ cap role is associated with the Xrn1-dependent buffering of the cellular mRNA levels, along with protecting from 5’-3’ degradation.

## INTRODUCTION

The N7-methylated guanosine (m7G cap) linked to the first nucleotide of the RNA molecule, is a hallmark structure of eukaryotic transcripts generated by the RNA Polymerase II (Pol II). The cap is involved in various steps of mRNA turnover, processing, transport and translation and therefore, essential for cell viability (Schwer, Mao and Shuman, 1998; Furuichi, 2015). Many of these biological functions are related to the cap-binding complex (CBC) which binds to the m7G caps of nascent RNAs (Izaurralde et al., 1995; Izaurralde et al., 1994).

The biosynthesis of the m7G cap requires three enzymes: the RNA triphosphatase (TPase) which removes the γ-phosphate from the RNA 5′ end; the RNA guanylyltransferase (GTase) which transfers a GMP group to the diphosphate 5′ end; and the guanine-N7 methyltransferase adding a methyl group to the N7 amine of the guanine cap (Decroly *et al.*, 2012). This synthesis pathway of the m7G cap is conserved in eukaryotes, however, the enzymes involved vary between the organisms (Ramanathan, Robb and Chan, 2016). In humans and other higher eukaryotes, a bifunctional protein RNGTT acts as RNA guanylyltransferase and 5′-triphosphatase while the methylation is mediated by a separate protein RNMT (Varshney *et al.*, 2018). In *S. cerevisiae* there are three proteins responsible for the RNA capping: the guanylyltransferase Ceg1 and the 5′-triphosphatase Cet1 work together as a stable heterodimer or heterotetramer, while the methyltransferase Abd1 acts independently (Schroeder *et al.*, 2000; Hausmann, Pei and Shuman, 2003; Martinez-Rucobo *et al.*, 2015). The capping enzymes are recruited to the Pol II at the early stage of transcription. In *S. cerevisiae*, Cet1 and Ceg1 bind the serine 5 phosphorylated C-terminal domain (CTD) of Pol II at the transcription start site (TSS) (Schroeder *et al.*, 2000; Lidschreiber, Leike and Cramer, 2013). When the nascent RNA reaches the length of ~17 nt, the Ceg1-Cet1 complex docks to the Pol II surface close to the RNA exit tunnel (Martinez-Rucobo *et al.*, 2015) and adds the terminal guanosine as soon as the first 25–30 nt of the nascent transcript extrude from the transcribing complex (Shatkin and Manley, 2000). Changes in the phosphorylation status of the Pol II CTD promoting transcription elongation results in the rapid dissociation of Ceg1-Cet1 (Martinez-Rucobo *et al.*, 2015). This and the increased phosphorylation of CTD Ser2 promote the binding of Abd1, which is recruited about 110 nt downstream of the TSS (Lidschreiber, Leike and Cramer, 2013) and completes the formation of the m7G cap. If the methylation of the added 5’ guanosine residue fails, the defective transcripts are detected by a surveillance mechanism that triggers decapping by Rai1 and subsequent RNA degradation by its co-factor, the 5’-3’ exonuclease Rat1 (Jiao et al., 2010). The lack of the cap structure on the 5’ end may also lead to co-transcriptional degradation of the nascent transcript. This process naturally occurs at the 3’ ends of genes and is essential for transcription termination. The cleavage over the poly(A) signal in the RNA creates an unprotected 5’ monophosphate RNA end in the nascent RNA. This serves as an entry point for Rat1 which co-transcriptionally degrades RNA pursuing the Pol II transcribing downstream of the gene (Kim *et al.*, 2004; West, Gromak and Proudfoot, 2004; Eaton *et al.*, 2020). If the degradation rate exceeds the RNA synthesis rate, Rat1 reaches the transcribing complex and Pol II is displaced from DNA (Cortazar et al., 2019; Eaton, Francis, Davidson, & West, 2020; Kim et al., 2004; West, Gromak, & Proudfoot, 2004). A similar mechanism has been suggested to be active upstream to the transcription end site (TES) when the capping fails at the 5’ ends of the genes. Analysis of the *FMP27* gene in *S. cerevisiae* revealed that the transcription of uncapped pre-mRNA is prematurely terminated by Rat1 within the gene body (Jimeno-González 2010). Moreover, analysis of the temperature-sensitive mutants revealed that the inactivation of Ceg1 affects the accumulation of heat-shock induced *SSA1* and *SSA4* mRNAs. However, their levels were restored by the deletion of the predominantly cytoplasmic 5’-3’ exonuclease Xrn1 (Schwer, Mao and Shuman, 1998). Xrn1 mediates the global mRNA degradation and therefore, together with the decapping factors, acts as a sensor of the cellular mRNA levels that controls and regulates the homeostatic mRNA abundance (Braun et al., 2012; Sun et al., 2013).

Here, we investigated the consequences of defective capping on RNA abundance and transcription. We took advantage of the yeast model where, in contrast to metazoans, CBC is not essential (Fortes *et al.*, 1999) which presents a unique opportunity to distinguish between the functions of m7G cap and CBC. We employed rapid nuclear depletion of Ceg1 to investigate the global turnover of the non-capped mRNAs. We found that the absence of the capping complex resulted in the differential expression of cellular RNAs: the mRNA levels of highly expressed genes decreased while the mRNAs of genes expressed at lower levels increased. The deletion of *XRN1* resulted in a similar redistribution in the mRNA abundance which was not synergistically affected by the nuclear depletion of Ceg1. Our data indicate that both, m7G cap and Xrn1 act closely in the control of the mRNA abundance. The non-capped mRNA levels were not rescued by the inactivation of Rat1 exonuclease and the disruption of the 5’ end capping had a minor effect on the distribution of the Pol II on the chromatin. Overall, our data indicate that the non-capped mRNAs are not simply evenly degraded and that the cap plays an important role in the Xrn1-dependent sensing and buffering of mRNA levels.

## RESULTS

### Differential expression of non-capped mRNAs

The impact of defective capping on the RNA expression has never been studied genome-wide before. In our experimental approach, we employed the anchor-away (AA) system (Haruki, Nishikawa and Laemmli, 2008) to deplete the capping enzymes from the nucleus and rapidly affect the formation of the cap. FRB tag fused with Cet1, Ceg1 or Abd1 (*cet1-AA, ceg1-AA* and *abd1-AA* strains) in the presence of rapamycin heterodimerizes with FKBP12 attached to the ribosomal protein Rpl13a, which is exported to the cytoplasm during the ribosome biogenesis, and thus removes the capping enzymes from the nucleus. Since rapamycin affects the TOR signalling pathway (Hausch et al. 2013) the strain used for this system carries the *tor1-1* mutation which permits normal growth on rapamycin (Haruki, Nishikawa and Laemmli, 2008). The nuclear depletion of Cet1 and Ceg1 were lethal (Fig. 1A) indicating the efficacy of the method as cap formation is essential for cell viability (Tsukamoto *et al.*, 1997; Chu and Shatkin, 2008). The inactivation of Abd1 activity has been shown to be lethal (Mao, Schwer and Shuman, 1996), however, the nuclear depletion of Abd1 had no impact on cell survival and only slightly affected the abundance of *ADH1* mRNA (Fig. S1A). This indicates that Abd1 may be still functional after the relocation to the cytoplasm. The nuclear depletion of either Cbp20 or Cbp80 was viable, consistent with the fact that the CBC is not essential in *S. cerevisiae* (Fig. 1A).

**Figure 1.**
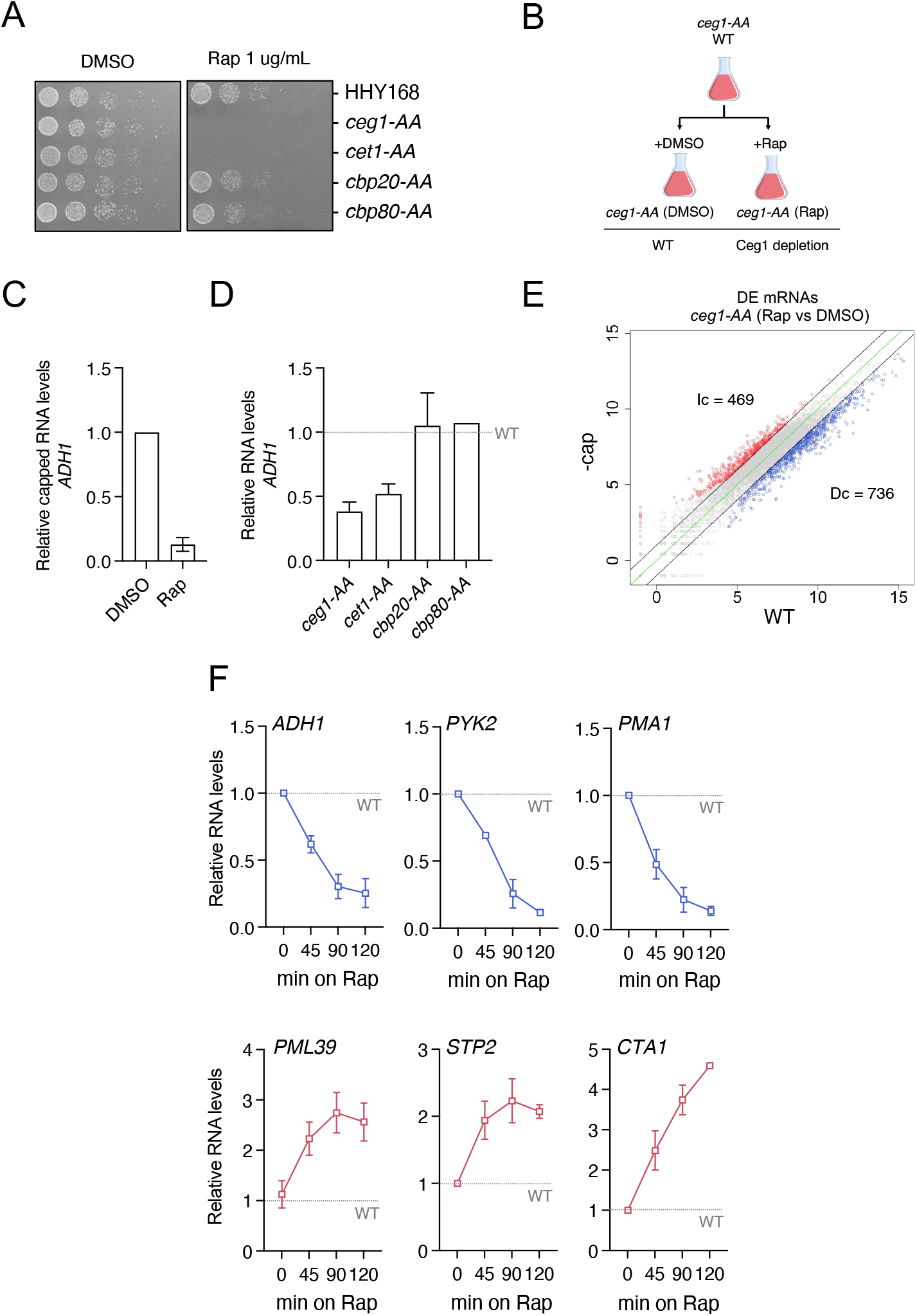
Nuclear depletion of capping enzymes results in differential accumulation of mRNAs. **A.** Spot test showing the growth of the AA strains. Serial dilution of each strain was spotted on solid YPD media in the presence of DMSO or rapamycin at the final concentration of 1 ug/mL. **B.** The outline of the experimental approach. Liquid cultures of cells at the exponential phase were split and supplemented with DMSO (control sample) or rapamycin (Rap; nuclear depletion sample) shown on the *ceg1-AA* example. **C.** Nuclear depletion of Ceg1 affects the formation of the m7G cap. Quantification of the capped RNA by RT-qPCR after 45 min of rapamycin treatment compared to control (DMSO). The RNA isolated from yeast was spiked-in with capped human RNA and immunoprecipitated by the anti-m7G cap antibody (clone H20). The RNA levels were normalised to human *GAPDH* mRNA. The error bars show standard deviation of three independent experiments. **D.** Nuclear depletion of the Cap Binding Complex does not affect mRNA levels. RT-qPCR analysis of the *ADH1* mRNA levels after 45 min of rapamycin treatment compared to the control set to 1 (dotted line). The error bars show standard deviation of two (*cbp20-AA* and *cbp80-AA*) or three independent experiments **E.** RNA-seq analysis showing differential expression of mRNA species in *ceg1-AA* (Rap) strain (-cap) after 45 min of rapamycin treatment compared to the control condition (WT). Increased (Ic) RNA species with log2 fold change >1 are labelled in red. Decreased (Dc) RNA species showing log2 fold change <1 are in blue. The zero-change line is in green and the ±1 log2 fold change threshold is indicated with continuous black lines. **F.** The changes in mRNA levels over time during nuclear depletion of Ceg1. RT-qPCR analysis of RNA levels for different genes measured at 0, 45, 90 and 120 min of rapamycin treatment relative to control (WT) set to 1 (dotted line). The error bars show standard deviation of three independent experiments.

In our functional experiments, the same yeast culture was split into a medium containing either DMSO or rapamycin representing the WT (DMSO) or the nuclear depletion (Rap) condition (Fig. 1B). First, we confirmed that RNA capping was indeed affected by the nuclear depletion of Ceg1 (Fig. 1C). We co-immunoprecipitated the RNA isolated from the *ceg1-AA* strain growing on either DMSO or Rap with the anti-m7G cap antibody (H-20). The amount of capped *ADH1* mRNA (normalised to human *GAPDH* spike) was reduced by 80% in *ceg1-AA* (Rap) when compared to WT (control cells grown on DMSO). The nuclear depletion of either Ceg1 or Cet1 (*ceg1-AA* or *cet1-AA)* resulted in a similar decrease in the *ADH1* mRNA (Fig. 1D and Fig. S1B) and therefore, in the subsequent analyses, we focused mainly on the effects of Ceg1 removal. We also confirmed that the nuclear depletion of CBC did not affect *ADH1* mRNA (Fig. 1D and Fig. S1B).

Next, we performed a global analysis to investigate genome-wide the mRNA levels after Ceg1 nuclear depletion. We depleted Ceg1 for 45 minutes (min) as this time was sufficient to decrease the *ADH1* level (Fig. S1B, 1D). Our RNA-seq analysis identified 1205 mRNAs affected by the nuclear depletion of Ceg1 (displaying log2 fold change < −1 and > 1) (Fig. 1E). 736 (61%) of these mRNAs decreased when compared to the control (WT), reflecting the fraction of non-capped transcripts that were subjected to degradation. Unexpectedly, the levels of 469 genes (39%) increased (Fig. 1E). Both effects were on average similar to each other (log2 fold change −1.4 and +1.5, respectively) (Fig. S1C). We also tested how the levels of the mRNAs changed over time during the nuclear depletion of Ceg1 (Fig. 1F). We selected *ADH1, PYK2* and *PMA1* genes whose levels decreased upon Ceg1 depletion in the RNA-seq data. RT-qPCR analysis revealed that their mRNAs decreased even further after 90 min and stabilised after 120 min. We also tested *PML39, STP2* and *CTA1* whose levels increased upon Ceg1 depletion in the RNA-seq data. *PML39* and *STP2* mRNAs peaked after 90 min, while the accumulation of *CTA1*, displayed a constant increase through time.

### Accumulation of non-capped mRNAs depends on their expression level

To understand the differential expression observed upon depletion of the capping machinery we analysed the features of the affected mRNAs. First, we tested the basal expression in WT cells (*ceg1-AA* on DMSO) for the-differentially expressed mRNAs in Ceg1-depleted cells. We found that increased mRNAs were generally lower expressed (median transcript per million, log2 TPM = 2.7) than the decreased mRNAs (median log2 TPM = 6.2) in WT (Fig. 2A). Next, we split all 1205 protein-coding genes differentially expressed in *ceg1-AA* into three bins according to their expression levels in WT cells: the top 25% were classified as highly expressed (High), the bottom 25% as lowly expressed (Low) and the remaining 50% as medially expressed (Mid) (Fig. 2B). Most of the lowly expressed genes increased their mRNA levels (291 out of 301), while most of the highly expressed mRNAs were decreased (300 out of 301) after the Ceg1 depletion. Mid-range expressed genes were the least affected by the depletion; however, they displayed the tendency to decrease the levels in Ceg1-depleted cells (426 decreased versus 176 increased), correlating with increasing expression (Fig. 2B-C). Overall, we observed a progressive decrease of the log2 fold change for mRNAs in *ceg1-AA* corelated with their increase in the basal expression levels in WT. This was also reflected at the single gene level as shown for *ZRT1, STB4* and *ADH1* (Fig. 2D). Next, we employed available datasets estimating mRNAs half-life (Geisberg et al., 2015), transcription rates (TR) and Pol II density in *S. cerevisiae* (Pelechano, Wei and Steinmetz, 2013) and applied them to the sets of differentially expressed mRNAs identified in the *ceg1-AA.* These analyses revealed that the mRNAs which increased upon Ceg1 nuclear depletion are more stable as they display longer half-lives (average half-life 41.9 vs 30.7 min for decreased mRNAs) (Fig. 2E and S2A), they also have lower transcription rates (Fig. 2F and S2B) and Pol II density (Fig. S2C) than the genes whose mRNAs decreased in Ceg1-depleted cells. The mRNAs accumulating in *ceg1-AA* were also longer than those from the decreased set (average gene length 1430 vs 1113 nt) (Fig. S2D) consistently with the fact that shorter genes are generally associated with higher expression (Urrutia and Hurst, 2003). Interestingly, the MEME suite (https://meme-suite.org/meme/) analysis identified a very pronounced sequence motif (A/G)GAAAA strongly enriched in the upregulated mRNAs (Fig. 2G) however, its function remains unknown.

**Figure 2.**
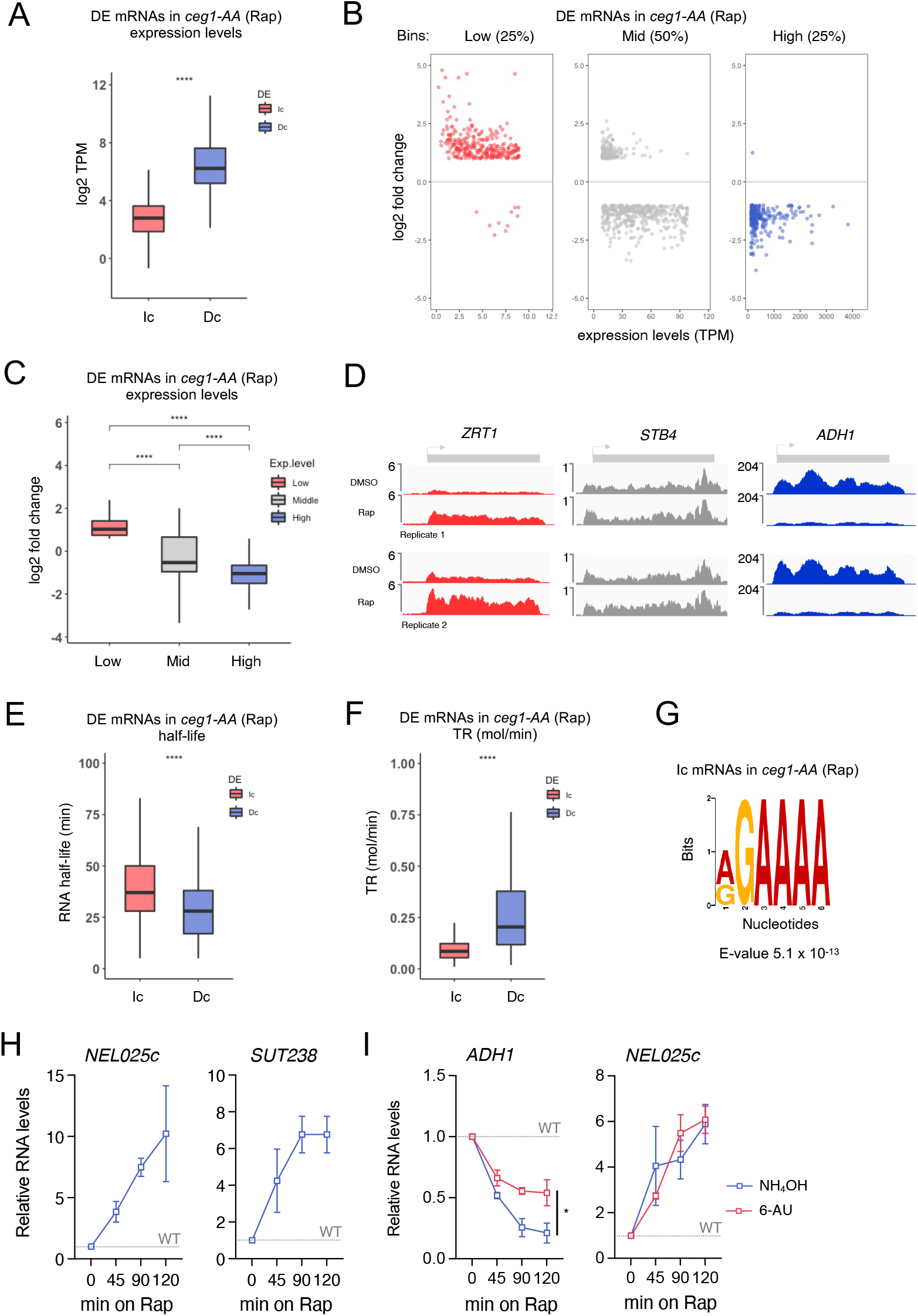
The accumulation of non-capped RNAs depend on gene expression levels. **A.** Differentially expressed mRNAs in *ceg1-AA* (Rap) vs *ceg1-AA* (DMSO/WT) sorted by their expression levels: Increased (Ic) or decreased (Dc). The expression levels were calculated by the log2 of Transcript Per Million (log2 TPM) value in WT (*ceg1-AA* on DMSO). **B.** The distribution of differentially expressed mRNAs in *ceg1-AA* (Rap) depending on their basal expression levels. **C.** The overall fold change for differentially expressed mRNAs in *ceg1-AA* (Rap) vs *ceg1-AA* (DMSO) ranked accordingly to their basal expression levels. **D.** Genome viewer tracks showing RNA-seq reads in *ceg1-AA* (DMSO) and *ceg1-AA* (Rap) for lowly (red), medial (grey) and highly (blue) expressed genes. **E-F.** Box plots showing half-life (E) and transcription rates (F) of differentially expressed mRNAs in *ceg1-AA* (Rap) vs *ceg1-AA* (DMSO). **G.** Logo sequence overrepresented in the increased mRNAs in *ceg1-AA* (Rap). **H.** The accumulation of the *NEL025c* (left) and *SUT238* (right) RNAs in *ceg1-AA* strain upon 0, 45, 90 and 120 min of rapamycin treatment compared to WT set to 1 (dotted line). **I.** RT-qPCR analysis showing the levels of *ADH1* mRNA and *NEL025* ncRNA upon 6-AU (red) and NH4OH (blue) treatments in Ceg1-depleted cells. The error bars show the standard deviation of three independent experiments. Statistical analysis is shown for the last timepoint (120 min). **A, C, E, F.** ns=P > 0.05; *=P ≤ 0.05; **=P ≤ 0.01; ***= P ≤ 0.001; ****= P ≤ 0.0001.

Gene ontology (GO) analysis showed that some mRNAs upregulated in *ceg1-AA* (45 out of the 469 tested) were associated with genes coding for DNA binding proteins directly associated with transcription (Fig. S3A) including the zinc finger protein *PPR1*,the regulator of stress response *STB5* and *PCF11*, a component of the 3’ end processing factor (Fig. S3B). GO terms analysis did not reveal consistent terms emerging from the genes whose mRNAs decreased in Ceg1-depleted cells (Fig. S3C).

We also investigated Pol II-transcribed non-coding RNAs (ncRNAs), called cryptic unstable transcripts (CUTs) and stable unannotated transcripts (SUTs), which in normal conditions are expressed at a very low level (Wyers *et al.*, 2005; Marquardt, Hazelbaker and Buratowski, 2011). Most CUTs are detectable only upon the inactivation of the 3’-5’ exonuclease Rrp6 subunit of the nuclear exosome (Xu *et al.*, 2009) and for this reason, only a handful of CUTs were detected in our RNA-seq analysis of the *ceg1-AA* strain. Thus, we focused our investigation on the well-studied ncRNA *CUT542* (*NEL025c*). The nuclear depletion of Ceg1 increased the levels of *NEL025c* (Fig. 2H) up to 10 times after 120 min of rapamycin treatment, consistent with the fact that lowly expressed Pol II transcripts accumulate when capping enzymes are not functional. A similar effect was observed for *SUT238* (Fig. 2H).

Since the expression of non-caped mRNAs depends on their basal expression, we tested whether the decrease in the RNA synthesis could alter the levels of *ADH1* mRNA and *NEL025c* ncRNA observed upon Ceg1 nuclear depletion. To affect Pol II elongation rate and processivity (Exinger and Lacroute 1992; Mason and Struhl 2005), we treated cells with 6-azauracil (6-AU) for 30 min, which causes depletion of intracellular nucleotide pools (Reines et al., 2003). This was followed by Ceg1 nuclear depletion and analysis of the *ADH1* and *NEL025c* levels relative to the WT (Fig. 2I). Slower RNA synthesis stabilised *ADH1* mRNA however, did not affect the overexpression of *NEL025c* which suggests that the low expression of this gene reached the threshold where further decrease had minimal impact on its accumulation.

### Degradation of non-capped Pol II transcripts

Previous reports suggested that the decreased mRNA levels caused by capping defects can be rescued by mutation of Rat1 exonuclease (Jimeno-González *et al.*, 2010). We employed *ceg1-63* and *ceg1-63/rat1-1* temperature-sensitive mutants to assess the levels of highly expressed *ADH1* as well as *FMP27* genes previously used to study capping-dependent RNA degradation (Jimeno-González *et al.*, 2010). *FMP27* is a lowly expressed gene and therefore is placed under the *GAL1* promoter in the *ceg1-63* background. We shifted the mutants and the parental strain to a non-permissive temperature (37 °C) up to 120 min (for the analysis of *GAL1::FMP27* the strain grew on galactose-containing medium). The mRNA levels of both *ADH1* and *FMP27* decreased over time in *ceg1-63* however, the Rat1 inactivation in *ceg1-63/rat1-1* did not rescue their abundance (Fig. 3A). The profile of the *FMP27* levels expressed from the *GAL1* promoter were mirrored by the levels of endogenous *GAL1* mRNA (Fig. 3A). In contrast to the temperature sensitive mutants *ceg1-63* and *ceg1-63/rat1-1*, native *FMP27* mRNA expressed at low level did not decrease in *ceg1-AA* but rather it remains stable (Fig. 3B). This may be caused by the difference in using ts mutants versus rapid nuclear depletion to induce the capping defects as well as by a significant difference between the expression from the native *FMP27* and *GAL1* promoters. Moreover, expression from the latter was already affected in the WT cells shifted to 37 °C. However, we cannot exclude that *FMP27* mRNA in *ceg1-63* is rescued by Rat1 inactivation at the sub-permissive temperature (34 °C) as previously described (Jimeno-González *et al.*, 2010). The RNA levels of lowly transcribed ncRNA *NEL025c* were not affected by the *rat1-1* mutation and increased in *ceg1-63/rat1-1* (Fig. 3A).

**Figure 3.**
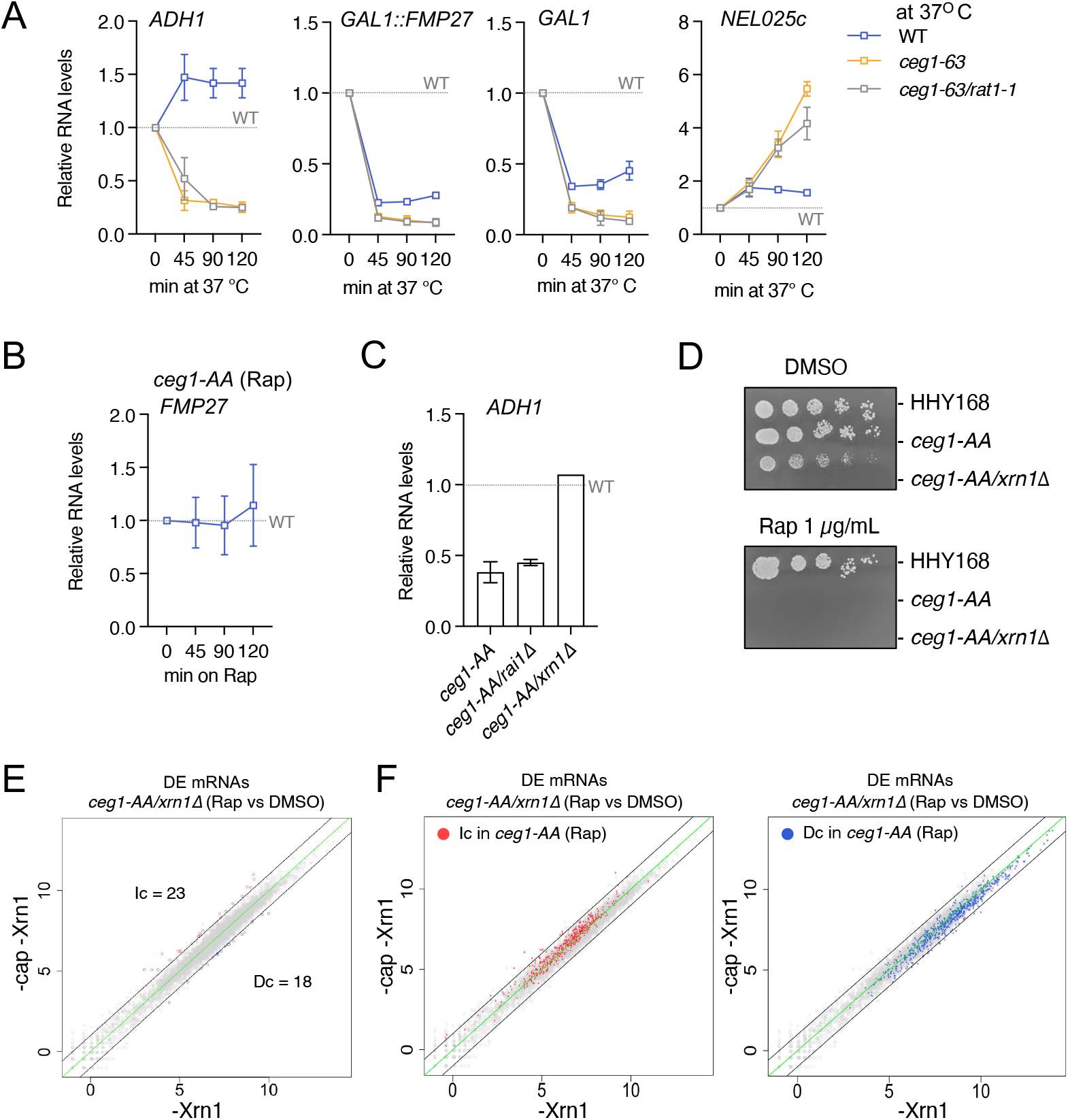
Roles for 5’3’ exonucleases in degradation of non-capped transcripts. **A.** RNA levels of *ADH1, FMP27, GAL1* and *NEL025c* in WT and temperature-sensitive mutants *ceg1-63* and *ceg1-63/rat1-1* after 0, 45, 90 and 120 min at a non-permissive temperature (37° C) compared to control (WT at 25° C) set to 1 (dotted lines). For analysis of *GAL1:FMP27* and *GAL1* mRNAs cells grew on gal-containing medium. **B.** *FMP27* mRNA levels in *ceg1-AA* after 0, 45, 90 and 120 min of rapamycin treatment compared to WT *(ceg1-AA* on DMSO) set to 1 (dotted line). **C.** *ADH1* mRNA levels in *ceg1-AA, ceg1-AA/rai1Δ* and *ceg1-AA/xrn1Δ* after 45 min on Rap compared to isogenic controls growing on DMSO set to 1 (dotted line). **D.** Spot test showing the growth of *ceg1-AA/xrn1Δ* on DMSO and rapamycin. Comparison with the growth of the parental strains *ceg1-AA* and HHY168. Serial dilution of each strain was spotted on solid YPD media in the presence of DMSO or rapamycin at the final concentration of 1 ug/mL. **E.** Differential expression of mRNAs after 45 min of rapamycin treatment in *ceg1-AA/xrn1*Δ (-cap -Xrn1) compared to the control condition *ceg1-AA/xrn1*Δ on DMSO (-Xrn1). Decreased (Dc) RNA species showing log2 fold change <1 are in blue. Increased (Ic) RNA species with log2 fold change >1 are labelled in red. The zero-change line is in green. The black lines represent the ±1 log2 fold change threshold. **F.** Increased (Ic) or decreased (Dc) mRNAs in *ceg1-AA* (Rap) are labelled on the plot showing mRNA levels in *ceg1-AA/xrn1*Δ (Rap) compared to the control condition *ceg1-AA/xrn1*Δ (DMSO). **A-C.** RT-qPCR. The error bars show the standard deviation of three independent experiments.

To avoid using the temperature-sensitive mutants we tested the Ceg1 nuclear depletion combined with the deletion of the genes encoding the Rat1 binding partner Rai1 (*ceg1-AA/rai1*Δ) or the exonuclease Xrn1 (*ceg1-AA/xrn1*Δ). The 5’ to 3’ exonucleolytic degradation mediated by Rat1 and Xrn1 requires a monophosphate RNA end (pRNA) while the RNA 5’ ends generated by Pol II contain three phosphate groups (pppRNAs) (Doamekpor et al., 2020; Jinek, Coyle, & Doudna, 2014). Triphosphorylated 5’ ends can be converted into proper substrates of the exonucleases by Rai1 in the nucleus. Rai1 also stabilises and activates Rat1 by forming a stable Rai1-Rat1 complex (Bresson and Tollervey 2018; Kachaev et al. 2020; Xiang et al. 2009). The Ceg1 nuclear depletion resulted in a similar decrease of *ADH1* mRNA in strains with either functional or deleted *RAI1* (Fig. 3C). This further confirmed that Rat1 is not efficiently involved in the degradation of non-capped transcripts. In contrast, the deletion of *XRN1* rescued the levels of the non-capped *ADH1* mRNA in *ceg1-AA* (Fig. 3C). Notably, the *ceg1-AA/xrn1*Δ strain grew slower than WT and was non-viable on Rap-containing medium (Fig. 3D), underlining the importance of m7G cap for cellular processes (e.g., translation).

The rescue effect observed for *ADH1* in *ceg1-AA/xrn1*Δ (Rap) prompted us to test whether the differential expression in *ceg1-AA* could be rescued globally by the deletion of the Xrn1 exonuclease. The *XRN1* null mutants were shown to stabilise non-capped mRNAs in *CEG1* temperature-sensitive mutants (Schwer, Mao and Shuman, 1998). We performed global differential expression analysis on the *ceg1-AA/xrn1*Δ strain growing on either DMSO (*xrn1*Δ) or Rap for 45 min. The Ceg1 nuclear depletion in the *xrn1*Δ background resulted in only 18 decreased mRNAs, 13 of which were also reduced in *ceg1-AA* (Rap) (Fig. 3E-F). Unexpectedly, the number of increased mRNAs was also significantly lower compared to *ceg1-AA* as we detected only 23 upregulated mRNAs (Fig 3E-F).

To elucidate why the *XRN1* deletion affected the accumulation of non-capped mRNAs in Ceg1-depleted cells, we tested mRNA expression in *ceg1-AA/xrn1*Δ prior to the nuclear depletion of Ceg1. We compared *ceg1-AA/xrn1*Δ (DMSO) to WT (*ceg1-AA* on DMSO) and found that *XRN1* deletion results in a global change in the mRNA expression, similar to the Ceg1 nuclear depletion. RNA-seq analysis revealed 543 upregulated and 605 downregulated mRNAs displaying log2 fold change < −1 and > 1 in *ceg1-AA/xrn1Δ* on DMSO (Fig. 4A). The upregulated genes were lowly expressed (median log2 TPM = 2.6) while the decreased fraction was highly expressed (median log2 TPM = 6.2) in WT cells (Fig. 4B). All of 1148 mRNAs differentially expressed in *xrn1*Δ were split into three bins according to their expression levels in WT cells (Fig. 4C-D). The majority of the lowly expressed mRNAs increased (270 out of 282), while most of the highly expressed mRNAs decreased (280 out of 282) in *xrn1Δ*. The number of mid-range expressed genes was equally affected (268 upregulated, 296 downregulated) however, the median of their expression levels was overall decreased. Consistently, upregulated genes displayed longer half-time compared to the downregulated with a median of 30 min and 17 min, respectively (Fig. 4E). Next, we tested if the effect of nuclear depletion of Ceg1 in *xrn1Δ* strain was masked by the pre-existing aberrant accumulation of mRNAs caused by the only *XRN1* deletion. We assessed how many differentially expressed mRNAs (log2 fold change < −1 and > 1) in *ceg1*-AA were also differentially expressed in *xrn1Δ* prior to nuclear depletion of Ceg1. We found that 190 out of 469 mRNAs increased and 264 out of 736 mRNAs decreased in *ceg1-AA* were also increased or downregulated respectively in *xrn1Δ* (Fig. 4F). This overlap identified only genes with log2 fold change over the ±1 threshold in both data sets therefore, we also tested the levels of differentially expressed mRNAs in *ceg1-AA* in the total mRNA fraction in *xrn1Δ*. Generally, most of the increased and decreased mRNAs identified in *ceg1-AA* were similarly affected in *xrn1Δ* (Fig. 4G) displaying a mean log2 fold change of 1.28 for the increased and −1.01 for the decreased mRNAs (Fig. 4H). We also performed reverse analysis and tested the levels of increased and decreased mRNA in *xrn1*Δ (log2 fold change < −1 and > 1) on all mRNAs in *ceg1-AA* (Fig. 4I). Both groups are accordingly increased or decreased in *ceg1-AA* (Rap) by a mean log2 fold change of 1.2 and −1.1 respectively (Fig. 4J). This analysis indicates that both *XRN1* deletion and lack of capping deregulate mRNA levels through a common mechanism and therefore the Xrn1 and Ceg1 co-inactivation did not result in differentially expressed genes (Fig. 3E). Finally, we compared the RNA-seq dataset from *ceg1-AA* (Rap) and *ceg1AA/xrn1*Δ (Rap) to exclude the genes similarly affected by either *XRN1* deletion or Ceg1 depletion. We identified 482 up- and 536 downregulated (Fig. S4A), both groups are similarly expressed in WT cells (Fig. S4B).

**Figure 4.**
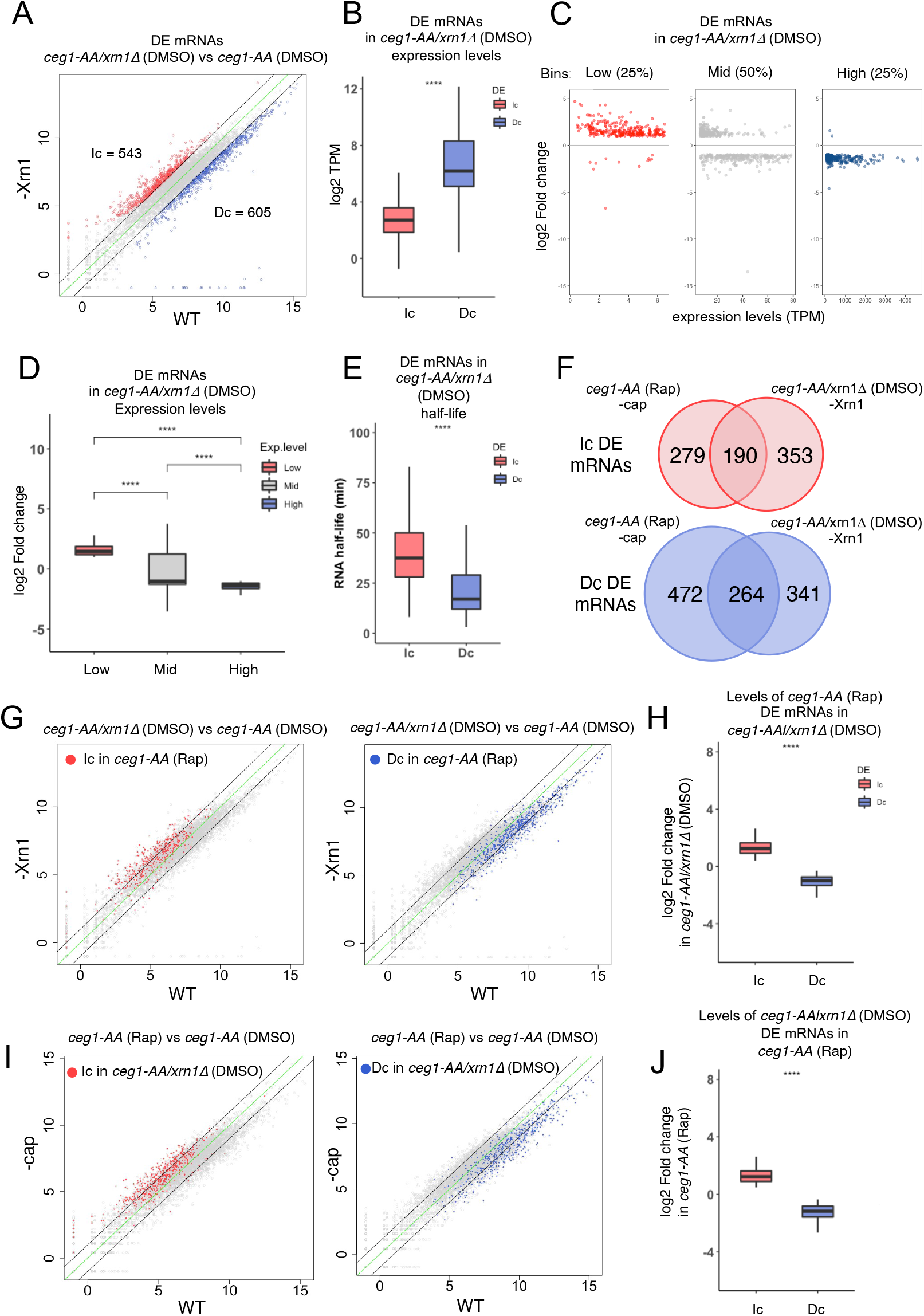
*XRN1* deletion and Ceg1 nuclear depletion similarly affect mRNA levels. **A.** Differential levels of mRNAs in *xrn1Δ* (*ceg1-AA/xrn1*Δ on DMSO) compared to WT (*ceg1-AA* on DMSO). Increased (Ic) mRNAs with log2 fold change >1 are labelled in red, decreased (Dc) mRNAs showing log2 fold change <1 are in blue. The zero-change line is in green. The black lines represent the ±1 log2 fold change threshold. **B.** Differentially expressed mRNAs in *xrn1*Δ (*ceg1-AA/xrn1*Δ on DMSO) sorted by their expression levels calculated by the log2 Transcript Per Million (log2 TPM) value in WT (*ceg1-AA* on DMSO). **C.** The distribution of differentially expressed mRNAs in *xrn1*Δ depending on their expression levels in WT cells. **D.** The overall fold change for differentially expressed mRNAs in *xrn1*Δ ranked accordingly to their expression levels in WT cells. **E.** The half-life of differentially expressed mRNAs in *xrn1Δ (ceg1-AA/xrn1Δ* on DMSO). **F.** The number of overlapping increased (Ic) or decreased (Dc) mRNAs in *xrn1*Δ (*ceg1-AA/xrn1*Δ on DMSO) and *ceg1-AA* (Rap). **G.** Increased (red) or decreased (blue) mRNA in *ceg1-AA* (Rap) are labelled on the plots showing mRNA levels in *ceg1-AA/xrn1*Δ (DMSO) compared to the control condition (WT). **H.** The log2 fold change in *xrn1Δ* for the sets of mRNAs that were differentially expressed in *ceg1-AA* (Rap). **I.** Increased (red) or decreased (blue) mRNAs in in *xrn1*Δ are labelled on the plots showing mRNA levels in *ceg1-AA* (Rap) compared to the control condition (WT). **H.** The log2 fold change in *ceg1-AA* (Rap) for the sets of mRNAs that were differentially expressed in *xrn1*Δ.

Our data indicate that non-capped mRNAs are not subjected to any specific quality-control degradation upon Ceg1 depletion, but they are rather degraded according to their expression. This and the similarities with the *xrn1*Δ mutation suggest that the cap structure is required for the processes of sensing and buffering the global mRNA levels by Xrn1. Non-capped mRNAs are not rapidly degraded upon Ceg1 depletion as they are not optimal substrates for the 5’-3’ exonucleases and require modifications of the 5’ ends by other enzymes. We hypothesise that such modifications are efficiently performed if the capping enzymes are in place which may represent a condition for the cap quality control to occur. In this scenario, even lowly expressed non-capped mRNAs with a suitable 5’ end would be efficiently removed in WT cells. To test this hypothesis, we introduced the self-cleaving hepatitis delta virus (HDV) ribozyme sequence into the 5’ UTR of the *FMP27* gene (*fmp27-RZ*) (Fig. 5A). The internal self-cleavage of *fmp27-RZ* generated a non-capped transcript with a 5’-OH end which is a suboptimal substrate for 5’-3’ exonucleases (Doamekpor et al., 2020; Jinek, Coyle, & Doudna, 2014). As a control we used the catalytically inactive ribozyme mutant harbouring the substitution C76:U (*fmp27-C76U*) (Perrotta, Shih and Been, 1999; Bird *et al.*, 2005). *FMP27* levels did not change upon Ceg1 nuclear depletion (Fig. 3B). However, in WT cells the introduction of the HDV ribozyme decreased the levels *of fmp27-RZ* by 64% compared to the catalytically inactive *fmp27-C76U* (Fig. 5B). This indicates that the generation of the correct substrate for exonucleases was required for the quality control and efficient degradation as previously reported (Bresson and Tollervey 2018).

**Figure 5.**
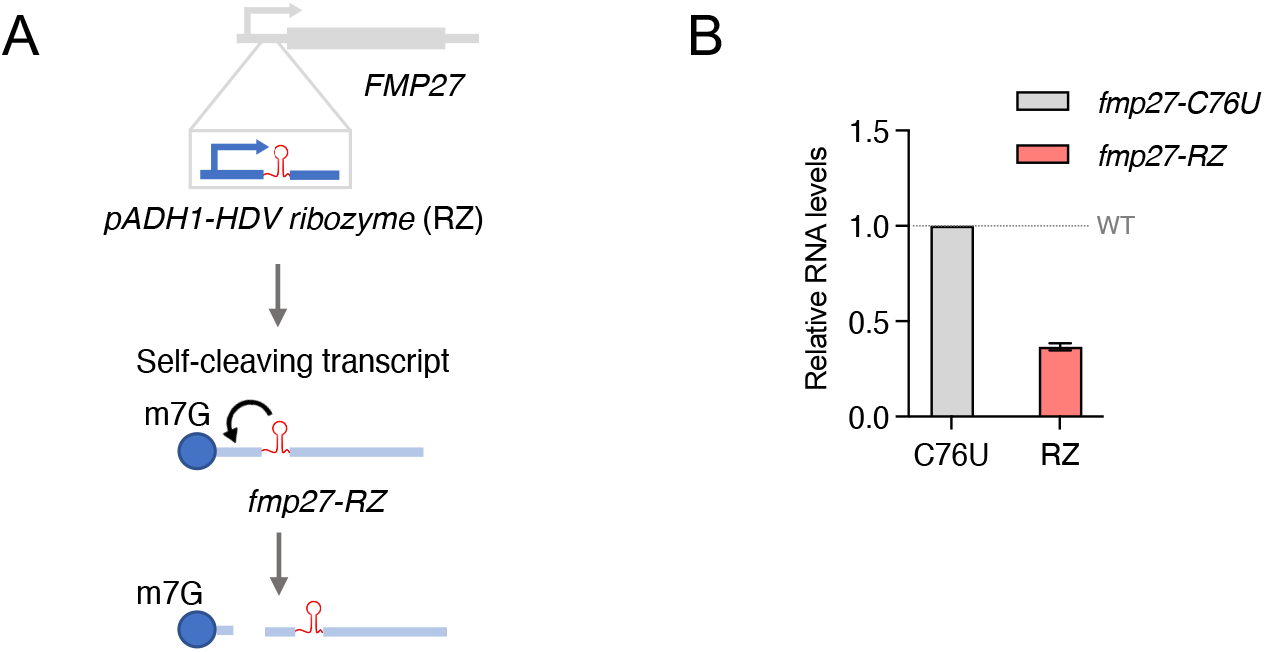
Cap removal is required for 5’-3’ degradation. **A.** The experimental approach for the generation of the self-cleaving *fmp27-RZ* transcript. **B.** The mRNA levels of *fmp27-RZ* and *fmp27-C76U* mutants. The error bars show the standard deviation of three independent RT-qPCR experiments.

### Non-capped mRNAs are not prematurely terminated

Although efficient co-transcriptional degradation is required for termination, previous studies reported that shifting the temperature-sensitive mutant *ceg1-63* to sub-permissive temperature resulted in a premature transcription termination of the 8 kb-long gene *FMP27* (Jimeno-González *et al.*, 2010). This effect was not fully recapitulated in the *ceg1-AA* strain. Our ChIP analysis using an antibody recognizing Pol II CTD revealed only a slight reduction of Pol II occupancy 0.5 kb from the transcription start site but not further downstream of the gene (Fig. 6A). Since the average gene length in yeast is 1.4 kb (Hurowitz and Brown, 2003), we investigated how transcription was affected by Ceg1 nuclear depletion on shorter genes (Fig 6B).

**Figure 6.**
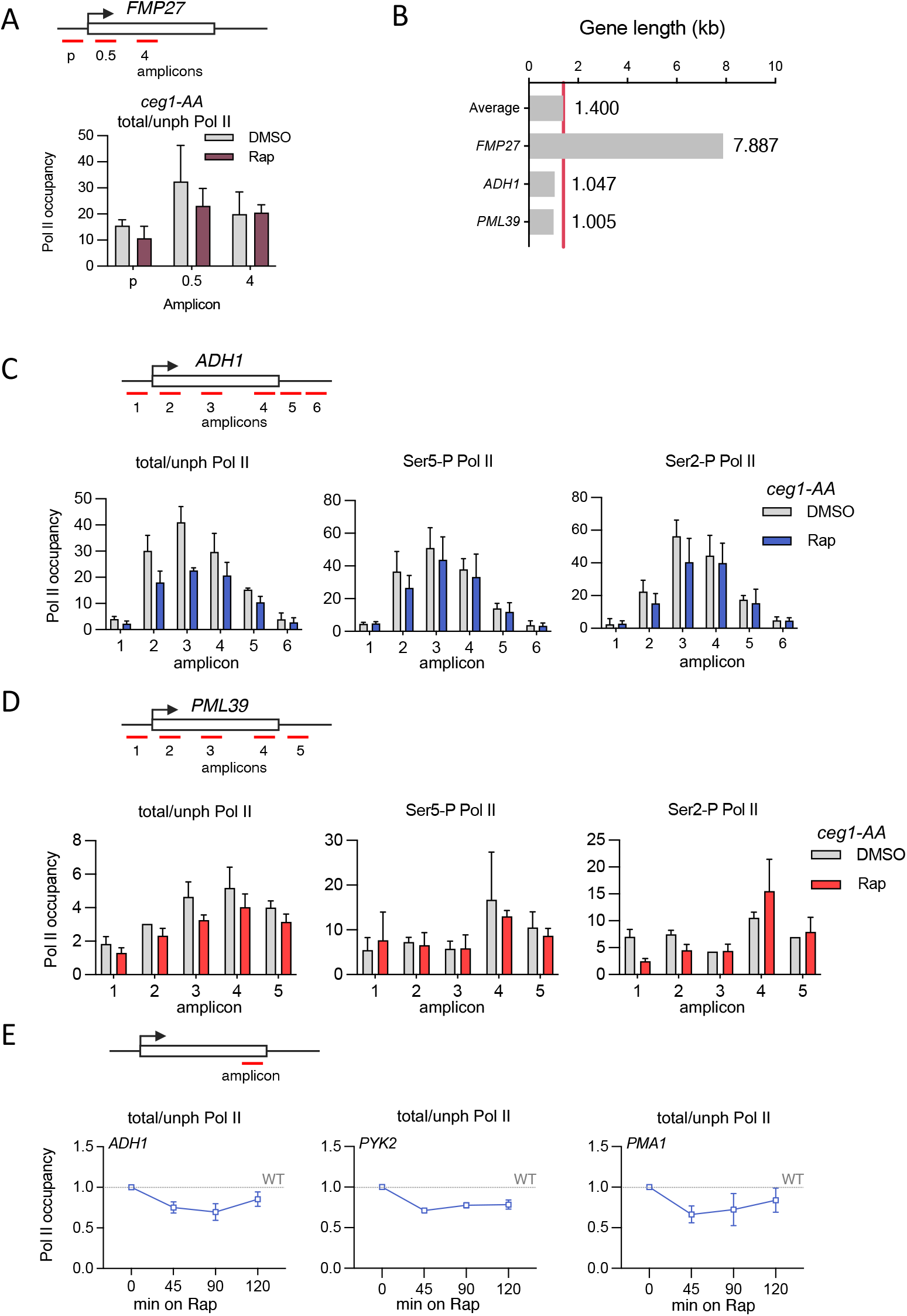
Transcription of non-capped mRNAs is not disrupted by pre-mature termination. **A.** The distribution of total/unphosphorylated Pol II (total/unph Pol II) over *FMP27* in control (DMSO) and after 45 min of rapamycin treatment (Rap) in *ceg1-AA*. The location of the amplicons used for the ChIP-qPCR over *FMP27* is shown above the chart. The error bars show the standard deviation of three independent ChIP-qPCR experiments. **B.** Chart showing the length of FMP27, *ADH1* and *PML39* genes. The red line represents the average gene length in *S. cerevisiae* (1.4 kb). **C-D.** The distribution of total/unphosphorylated Pol II (total/unph Pol II) as well as serine 5-phosphorylated (Ser5-P Pol II) and serine 2-phosphorylated (Ser2-P Pol II) Pol II isoforms over *ADH1 (C)* and *PML39* (D) in control (DMSO) and after 45 min of rapamycin (Rap) in *ceg1-AA* strain. The location of the amplicons used for the ChIP-qPCR is shown above the charts. The error bars show the standard deviation of three independent ChIP-qPCR experiments. **E.** The occupancy of the total/unphosphorylated Pol II at the 3’ end of *ADH1, PYK2* and *PMA.* Timepoints after 0, 45, 90 and 120 min on rapamycin compared to control (*ceg1-AA* on DMSO) and set to 1 (dotted line). The location of the amplicons used for the ChIP-qPCR is shown above the charts. The error bars show the standard deviation of three independent ChIP-qPCR experiments.

We depleted Ceg1 for 45 min and analysed the distribution of Pol II phosphoisoforms on highly and lowly expressed genes: *ADH1* and *PML39*, respectively (Fig. 6C-D). ChIP analysis revealed that total Pol II was partially depleted mainly on the 5’ ends for both highly and lowly expressed genes. We used 8WG16 antibody recognising non-modified heptapeptides of the CTD to analyse total Pol II distribution. Thus, we speculate that this decrease of the Pol II signal detected by this antibody may reflect prematurely terminated unphosphorylated or hypophosphorylated Pol II. Indeed, both ser5-P and ser2-P Pol II phosphoisoforms, that represent actively transcribing Pol II, were only marginally affected by Ceg1 depletion (Fig. 6C-D).

A similar but less pronounced effect was observed for *ADH1* gene in *cet1-AA* (Rap) (Fig. S4A) as both enzymes are essential for cap formation (Martinez-Rucobo *et al.*, 2015). Nuclear depletion of Abd1 did not affect the distribution of Pol II over *ADH1* (Fig. S4B). We also investigated if longer nuclear depletion of Ceg1 resulted in increased premature transcription termination. We shifted cells to the medium containing Rap for up to 120 min and tested the levels of Pol II over the 3’ end of high and mid-range expressed *ADH1, PYK1* and *PMA1* genes (Fig. 6E). Consequently, we observed only ~20% decrease in Pol II signal at 45 min which was not exacerbated by longer nuclear depletion of Ceg1.

## DISCUSSION

Capping is the first co-transcriptional RNA modification posed on the mRNAs. The m7G cap is essential for the functionality of the mRNA and therefore, mutations affecting the cap formation are non-viable. Previous studies reported that RNAs with defective caps (e.g. lacking m7G or with non-methylated caps) are rapidly degraded by cap quality control mechanisms, however, these analyses were performed only on a few selected genes (Lewis and Izaurralde, 1997; Schwer, Mao and Shuman, 1998; Cougot *et al.*, 2004; Topisirovic *et al.*, 2011). Here, we performed a global analysis of the capping defective transcriptome. We show that although the overall mRNA levels decrease was a dominant effect following the nuclear depletion of the capping enzymes, the levels of many mRNAs did not change and strikingly, some of the mRNAs increased when not capped. Our data show a clear dependence between the amount of RNA synthesised, stability and the degradation of uncapped transcripts. The nuclear depletion of Ceg1 resulted in the depletion of highly expressed mRNAs and accumulation of lowly expressed and more stable mRNAs. We observed the same pattern in cells lacking the 5’-3’ exonuclease Xrn1 which is the major enzyme degrading mRNA in the cytoplasm. Consistently, we did not detect any significant number of differentially expressed genes in the *xrn1D* strain following nuclear depletion of Ceg1. These observations allowed us to draw a few conclusions. First, our data indicate that non-capped mRNA is not quality controlled but randomly degraded. Therefore, the most abundant and so most accessible RNA species decrease upon nuclear depletion of the capping machinery. This in turn, may increase the relative concentration of lowly transcribed mRNAs in the overall fraction which we detected as upregulated mRNAs. We speculate that the m7G cap may not be essential for the protection from exonucleolytic 5’-3’ degradation upon Ceg1 depletion, as the 5’ triphosphate non-capped mRNAs synthesised are not immediate substrates of the 5’-3’ exonucleases. Cells have mechanisms that convert different variants of RNA 5’ ends to substrates for exonucleases. For example, Rai1 shows both decapping and pyrophosphohydrolase activity which generates monophosphate RNA 5’ end (pRNA) substrate of its binding partner, the 5’-3’ ribonucleases Rat1 (Xiang *et al.*, 2009; Jiao *et al.*, 2010). However, the depletion or inactivation of these proteins did not rescue the mRNA levels in Ceg1-depleted cells, indicating that the Rai1-Rat1 complex is not efficiently involved in their degradation. Similarly, the levels of nuclear CUT and SUT tested in this study increased upon simultaneous Ceg1 depletion and Rat1 inactivation. The lack of a specific degradation pathway targeting the RNAs synthesised without the cap may aid the production of other essential RNA classes produced by the RNA Pol I and III such as ribosomal RNA (rRNA) or transfer RNA (tRNA), respectively, that are not capped but stable in the nucleus and cytoplasm (Ramanathan, Robb and Chan, 2016; Akiyama, Eiler and Kieft, 2017). In these cases, the synthesis would not be affected by the surveillance mechanisms degrading non-capped RNAs in the nucleus. However, releasing and maintaining non-functional mRNAs may have deleterious effects on the cell. Thus, we speculate that the binding of the capping enzymes to the transcriptional machinery or the synthesis of the m7G cap itself, is the condition required for the cap quality control pathway to be functional. Indeed, the insertion of the self-cleaving ribozyme into the 5’ UTR of *FMP27* resulted in degradation of this lowly transcribed mRNA in WT cells.

Cap may be essential for the Xrn1-dependent buffering of the global mRNA levels. Decapping enzymes interact with Xrn1, and the cap removal is the first step triggering mRNA degradation (Schoenberg and Maquat, 2012). Thus, non-capped mRNAs may not be identified as coding transcripts and therefore subjected to level correction. The hypothesis that the cap is required for efficient mRNA degradation and buffering of the mRNA levels is supported by the observation of a similar pattern in the mRNA levels in the strain lacking Xrn1. Almost all genes upregulated and downregulated upon Ceg1 nuclear depletion were also respectively affected in the *xrn1*Δ strain. We speculate that the lack of the cap may disturb the correct buffering of mRNA levels (Fig. 7) and thus mRNAs are degraded randomly by cytoplasmic degradation enzymes like the exosome.

**Figure 7.**
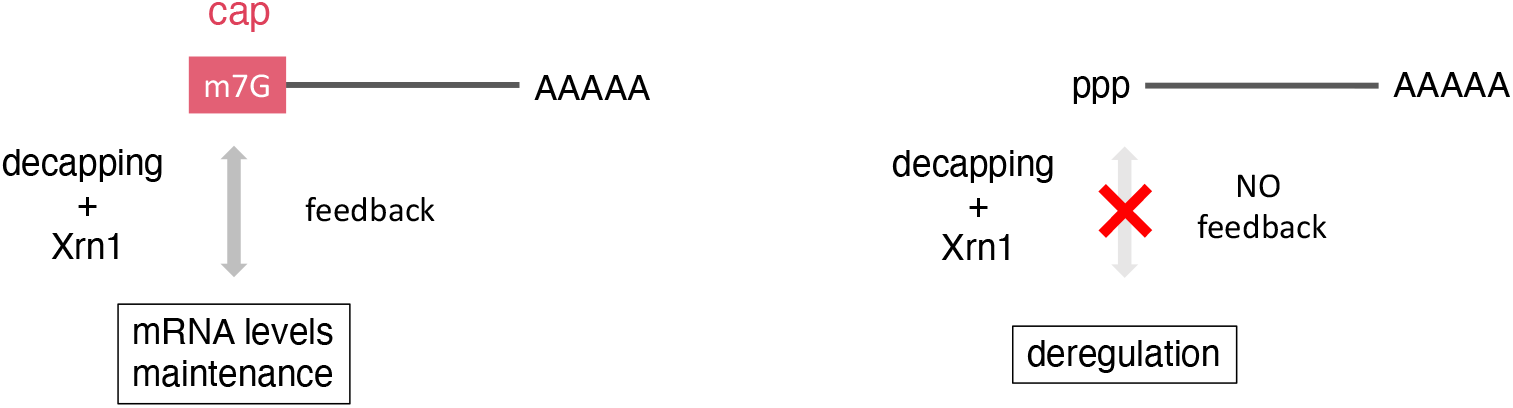
A model of cap-dependent regulation of mRNA levels.

Indeed, the cap plays a role in marking the Pol II-dependent transcripts. For example, the presence of the cap at the 5’ end decides the fate of yeast box C/D snoRNAs, a class of RNAs which do not possess the m7G cap as mature species. However, when their 5’ end processing is affected and the cap retained, these snoRNAs are subjected to mRNA processing pathways, become polyadenylated at the 3’ end as well as exported to the cytoplasm (Grzechnik *et al.*, 2018). Similarly, the cap structure may mark the mRNAs as coding transcripts facilitating the homeostatic adjustments in their levels by degradation and synthesis.

Finally, we did not observe a significant increase in premature transcription termination upon depletion of the capping enzymes. This is consistent with our data showing that Rat1 and Rai inactivation did not restore the levels of non-capped mRNAs, but in contrasts with previous models suggesting that such RNAs are degraded co-transcriptionally by Rat1 eliciting the removal of Pol II from DNA by the “torpedo termination” mechanism, where 5’-3’ exonuclease Rat1 attacks the 5’ unprotected end, chases the Pol II by degrading the nascent RNA, and ultimately displaces the transcribing polymerase from the DNA template (Eaton et al., 2020; Kim et al., 2004; West, Gromak, and Proudfoot, 2004). Our ChIP analysis revealed that only the fraction of Pol II tested using 8WG16 antibody was slightly depleted in *ceg1-AA* (Rap). This antibody recognises unphosphorylated heptapeptides in the CTD and therefore the precipitated fraction may be enriched for hypophosphorylated Pol II. Such Pol II may not efficiently transit to the elongation phase, pause or transcribe at a low speed (Bartkowiak and Greenleaf, 2011; Harlen *et al.*, 2016; Harlen and Churchman, 2017). All these conditions facilitate transcription termination by a torpedo mechanism, which may contribute to the lowered levels of Pol II observed in the ChIP analyses employing 8WG16 antibody. We speculate that for average length genes (1.4 kb) the window in which Rai-Rat1 can be recruited to the transcribing complex and thus catch normally transcribing Pol II is too narrow to trigger transcription termination upstream of the canonical transcription end site. Alternatively, the capping process must be at least initiated, and the capping enzymes present on the transcribing complex to facilitate the quality control mechanism. For example, premature transcription termination is observed when nascent RNAs are decapped which triggers Pol II pausing and consequent Rat1-dependent termination (Brannan *et al.*, 2012; Contreras, Benkirane and Kiernan, 2013; Eaton and West, 2020; Eaton *et al.*, 2020). Such condition would be consistent with our hypothesis that the major role of the cap is marking the mRNA as a product of the Pol II and coding RNAs in the cytoplasm.

Overall, our work indicates that the 5’ cap role is more complex than serving as a platform for the Cap Binding Complex and punctuating mature 5’ ends of Pol II transcripts. Cap may play an essential role in marking mRNA molecules and cooperating with Xrn1-dependent buffering of the cellular mRNA levels.

## ACKNOWLEDGMENTS

This work was funded by Sir Henry Dale Fellowship from the Wellcome Trust and the Royal Society (218537/Z/19/Z). O.Z. was supported by Midlands Doctoral Training Partnership by a studentship to O.Z. funded by BBSRC (grant number: BB/M01116X/1). DH is funded by EPSRC (grant EP/T002794/1). We thank Kinga Winczura, Lauren Garbett and Mark Ashe for critical reading of the manuscript and Genomics Birmingham at the University of Birmingham for assistance with NGS.

## CONFLICT OF INTEREST

The authors declare that they have no conflict of interest.

## AUTHOR CONTRIBUTION

O.Z. performed all experiments, bioinformatics and analysed data. D.H. supported bioinformatics. P.G. and O.Z. designed experimental approaches and wrote the manuscript.

## MATERIALS AND METHODS

### Data availability

The RNA-seq data for the *ceg1-AA* strain upon 45 min of treatment with DMSO or rapamycin are available in Gene Expression Omnibus (GEO) database with the accession number GSE213942.

### Yeast strain and Media

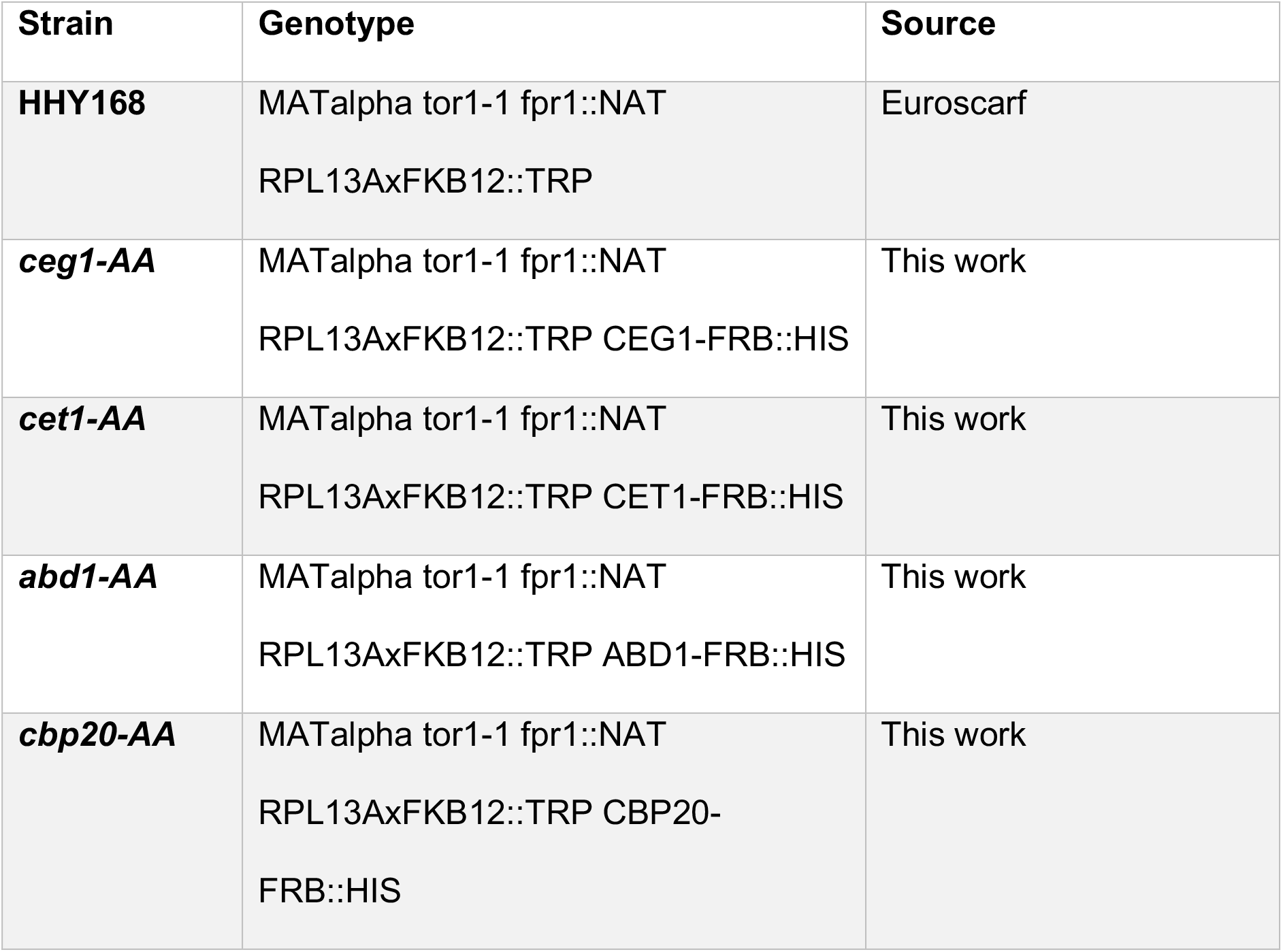
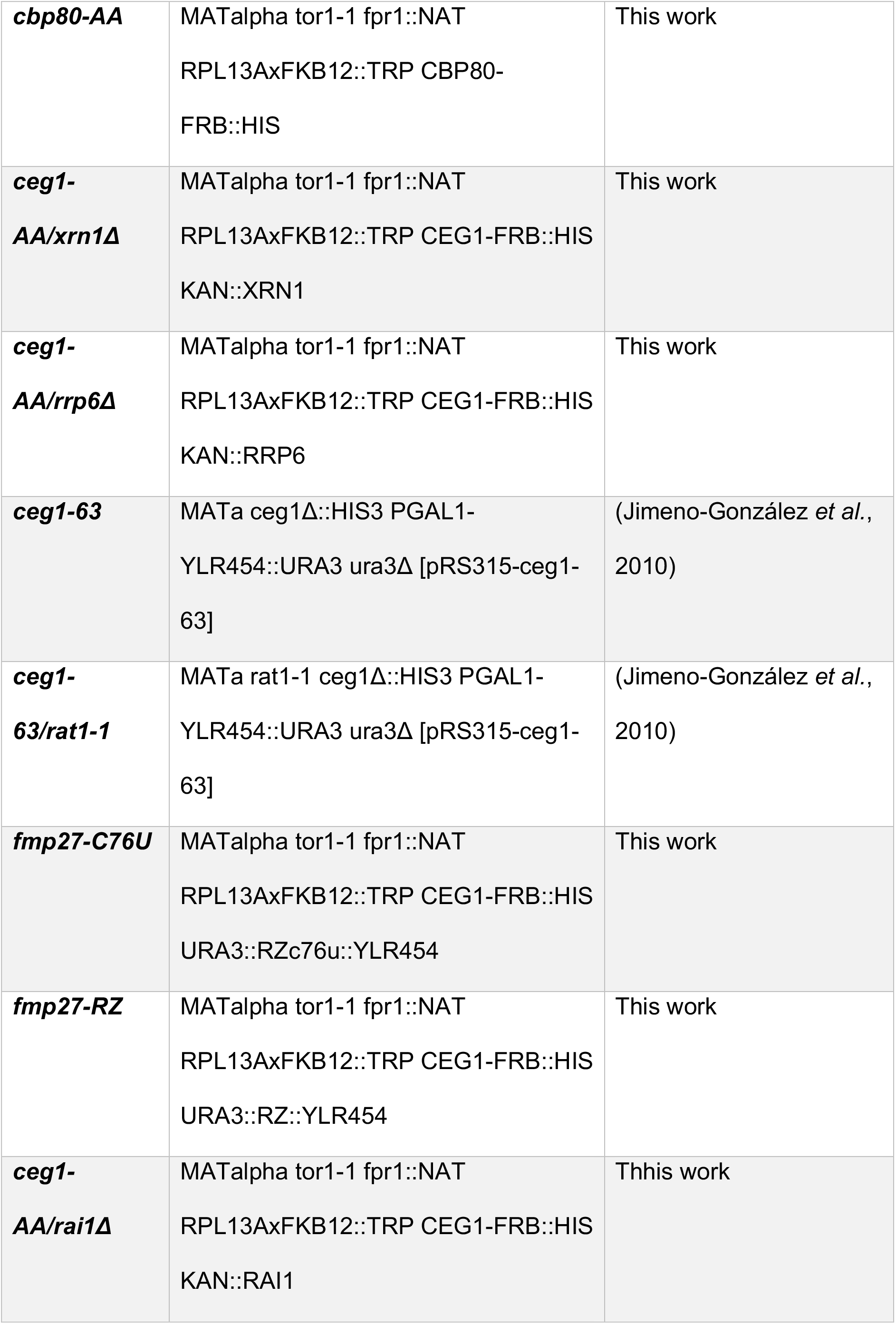

Yeast strains were grown at 30° C on YPD (1% yeast extract, 2% peptone, 2% dextrose/glucose) media enriched in DMSO or rapamycin to the final concentration of 1 μg/mL where indicated. For 6-AU treatment, the yeast cells were grown in minimal media (yeast nitrogen base without amino acids) with synthetic dropout ura-. For solid media, agar was added to the final concentration of 2%.

### Growth spot assay

Yeast cells were grown to OD600 = 0.4. Serial dilutions 1:25 were performed and 5 μL of each dilution spotted onto YPD (2% agar) plates.

### Yeast cell transformation

Cells were cultured to OD600 = 1 and pelleted by centrifugation for 5 min at 2 400 × g at room temperature (RT). The pellet was resuspended in sterile 10 mM Tris-HCl pH 7.5 or water and centrifuged again as before. The pellet was resuspended in filtered LiT (10 mM Tris-HCl pH 7.5;100 mM Lithium acetate), 1 M DTT and incubated at RT for 40 min with gentle shaking. After incubation the cells were pelleted again (as before) and resuspended in LiT and 1 M DTT. The suspension was added with LiT; and dsDNA (10 mg/mL); transforming DNA (0.1 to 1 ug or more) and incubated at RT for 10 min. Next, PEG solution (1:1 PEG4000 : LiT) was added to the suspension and incubated at RT for 10 min. DMSO was then added, and the suspension incubated at 42° C for 15 min. After the incubation the suspension was pelleted and resuspended in 1 mL of YPD (1% Bacto yeast extract, 2% Bacto peptone, 2% glucose/dextrose). Then the suspension was incubated at 30° C for 1 hour, pelleted with 10 seconds (sec) spin, max speed and inoculated on selective plates.

### RNA extraction

RNA was purified by phenol/chloroform method. The yeasts were cultured to the final concentration OD600 = 1, pelleted in 50 mL falcon tubes by centrifugation for 2 min at 1000 rpm at room temperature (RT). The pellet was frozen at −80° C before further manipulation. Next, the pellet was resuspended in 1 mL of ice-cold RNase free water, moved to a 1.5 mL tube, centrifuged for 10 sec at 4° C max speed. Then the supernatant (water) was removed, and the pellet resuspended in 400 μL of filtered AE buffer (50mM Sodium acetate pH5.3; 10mM EDTA), 40 μL of 10% Sodium dodecyl sulphate (SDS) and 400 μL of Acid Phenol (pH 4). The suspension was vortexed for 20 sec and incubated at 65° C for 10 min, then incubated for further 10 min at −80° C. The defrosted sample was then centrifuged for 5 min, 13 000 rpm at RT. 400 μL of 1:1 phenol:chloroform solution were added to the supernatant and vortexed for 30 sec. Then the sample was centrifuged 10 min, 13 000 rpm at RT. The upper phase was transferred and mixed with 400 μL of Chloroform, vortexed and centrifuged for 5 min, 13 000 rpm at RT. The RNA was precipitated by adding to the upper phase 1 mL of 100% ethanol and 40 μL of 7.5 M Ammonium acetate. Left at −80° C for 2 hours (hrs) then centrifuged for 20 min, 13 000 at 4° C. The pellet was washed with 70% ethanol (in water) and resuspended in RNase free water.

### RT-qPCR

Reverse transcription was performed with Super Script III Reverse transcriptase (Thermo Fisher, 18080044) following manufacturer’s instruction using random hexamers (Thermo Fisher, N8080127) for cDNA synthesis. qPCR data were analysed with the ΔΔCt method normalised to 25S ribosomal RNA (rRNA) and control condition.

### RNA sequencing

Before library preparation, ribosomal RNA (rRNA) depletion was performed on 5 μg of total yeast RNA using 200 pmol of 3’ biotinylated probes (IDT) designed in house following the RiboPOP method (Thompson, Kiourlappou, and Davis 2020). The libraries preparation was performed with Ultra II RNA Library Prep Kit for Illumina (NEB, E7770) and Multiplex Oligos for Illumina (NEB, E7335) according to manufacturer’s instructions. The RNA sequencing was performed from the Genomics Facility at the University of Birmingham on Illumina NEXTseq apparatus.

### RNA-seq data analysis

The quality of the data was checked using FastQC v0.11.5 (Andrews S. (2010). FastQC: a quality control tool for high throughput sequence data. Available online at: http://www.bioinformatics.babraham.ac.uk/projects/fastqc).

Reads were aligned to the yeast genome R64 (sacCer3) using HISAT2 v2.1.1 and sorted and indexed using SAMtools v1.15.1 (Wysoker et al. 2009). The read count per gene were calculated with LiBiNorm v.2.5 (Dyer, Shahrezaei and Hebenstreit, 2019) with the following parameters count -u name-root -f --order=pos --minaqual=10 -- mode=intersection-strict --idattr=gene_id --type=exon. Differential expression analysis was performed using DESeq2 package. GO analysis was generated using g:profiler (https://biit.cs.ut.ee/gprofiler).

### Cap Immunoprecipitation

Total RNA from cells treated with DMSO or rapamycin was extracted and use for cap pull-down. 5 μg of total yeast RNA was spiked with 1 μg of total human RNA as immunoprecipitation efficiency control. Capped RNA species were immunoprecipitated using 5 μg of anti m7G cap antibody (H-20) as previously described (Jimeno-González et al. 2010).

### 6-azauracil (6-AU) treatment

Overnight culture was prepared inoculating a single colony on minimal media as described above. The next day the culture was diluted to OD600 = 0.2. The diluted culture was grown with agitation at 30° C to OD600 = 1 and split in 4. Two cultures were added of 6-azauracil (6-AU) and the other two with ammonium hydroxide (NH4OH, control) at the final concentration of 50 μg/mL for 30 min. Next, DMSO or rapamycin (Rap) at the final concentration of 1 μg/mL were added according to the following scheme: 6-AU+DMSO, 6-AU+Rap, NH4OH+DMSO, NH4OH+Rap. 10 mL of each suspension were collected a time 0, 45, 90 and 120 min after the addition of DMSO or Rap and prepared for RNA purification.

### 4-Thiouracil (4tU) labelling and purification

Liquid yeast culture traded for 40 min with rapamycin (1 μg/mL) or DMSO were added of 4tU 500 μM final concentration (Sigma-Aldrich, 440736) to final concentration 5 mM for 5 min and subsequently flash frozen in cold ethanol (−80° C). The suspension was pelleted RNA purified by Phenol/Chloroform extraction. 200 μg of total RNA from *S. cerevisiae* was spiked with *S. pombe* total RNA (4:1 ratio *S. cereviasiae* : *S. pombe)*treated with 4-tU 500 μM final concentration (Sigma-Aldrich, 440736). The biotinylation was performed with 200ul EZ-link-HPDP-biotin (Thermo Fisher, 21341) for 30 min at 65° C. The RNA was then incubated with 5M sodium chloride and isopropanol and incubated with Dynabeads M-270 Streptavidin (Thermo Fisher, 65305) for 1 hour at room temperature. Beads were washed five times with Binding & Washing buffer (5 mM Tris-HCl pH 7.5, 0.5 mM EDTA and 1 M NaCl) and 4tU labelled-RNA was eluted with 100 μM DTT, precipitated with 5M NaCl, isopropanol and glycogen, and resuspended DNase/RNase free water. Enrichment of nascent RNA was confirmed with qPCR using exon–intron primers of ACT1 gene.

### Chromatin immunoprecipitation (ChIP)

The specific yeast strain was grown over-night to OD600 = 1. Then 37% formaldehyde was added to the media to final concentration 1% dropwise with gentle shaking for 15 min. Next, 15 mL of 2.5 M glycine was added for further 5 min keeping the shaking. The cells were pelleted by centrifugation at 3 500 rpm for 3 min at 4° C. The pellet was resuspended in 40 mL of ice-cold 1× PBS and centrifuged at 3 500 rpm for 5 min at 4° C, twice. The pellet obtained was then transferred and prepared for lysis. The pellet was resuspended in 1 mL of ice-cold FA1 (50 mM HEPES-KOH, pH7.5; 150 mM NaCl; 1 mM EDTA; 1% triton-x 100; 0.1% Sodium deoxycholate) added with proteinase inhibitor (Roche Mini-EDTA free) and about 400 μL zirconia beads. The cells were lysed in MagnaLyser (30 sec 7000 rpm, 3 cycles with 5 min incubation on ice in between). The lysate was drained out of the beads by centrifugation 1 000 rpm for 1 min at 4° C and SDS was added to a final concentration of 0.05%. The lysate was sonicated as follow: 25 cycles 15 sec ON, 15 sec OFF on maximum power. Next, it was centrifuged for 20 min at 12 500 rpm at 4° C. 200 μL of lysate was saved to check the fragmentation. SDS was added to the lysate with to final concentration of 1% and 0.05 mg of Proteinase K and incubated at 42° C for 1 hour. Next it was moved to 65° C for 4 hrs. The DNA de-crosslinked was precipitated by Phenol/Chloroform, the contaminant RNA removed by adding 2 μL of RNase A incubating for 1 hour at 37° C and checked on agarose gel. The rest of the lysate was centrifuged max speed for 5 min at 4° C. 150 μg of chromatin were used for immunoprecipitation. FA1 enriched with proteinase inhibitor was added to the DNA to a final volume of 700 μL. 20 μL of such mix was saved as input for the further ChIP analysis. 2 μg of the specific antibody was added to the sample and incubated on rotation over-night at 4° C.

15 μL of protein A (Thermo Fisher, 10002D) and 15 μL of protein G-coated beads (Thermo Fisher, 10004D) were washed with 1 mL of FA1 and resuspended in 100 μL of FA1. The beads were added to the sample and incubated by rotation for 1 hour at 4° C. After incubation the beads were washed 6 times with FA1, 1 time with FA2 (50 mM HEPES-KOH, pH 7.5; 500 mM NaCl; 1 mM EDTA; 1% Triton-X 100; 0.1% Sodium deoxycholate), 1 time with FA3 (20 mM Tris pH 8.0; 250 mM LiCl; 0.5% NP-40; 0.5% Sodium deoxycholate; 1 mM EDTA) and TE buffer pH 8 (100 mM Tris-HCl; 10 mM EDTA). The beads were then resuspended in 195 μL of Elution buffer (50 mM Tris pH 7.5; 10 mM EDTA; 1% SDS) and incubated for 10 min at 65° C and 0.05 mg of Proteinase K added. The Chromatin was then de-crosslinked by incubating the sample at 42 ° C for 1 hour and moved at 65° C for 4 hrs. The DNA was finally purified using Quiagen cleanup (28104) kit following the manufacturer’s instruction.

## SUPPLEMENTAL FIGURES

**Figure S1.**
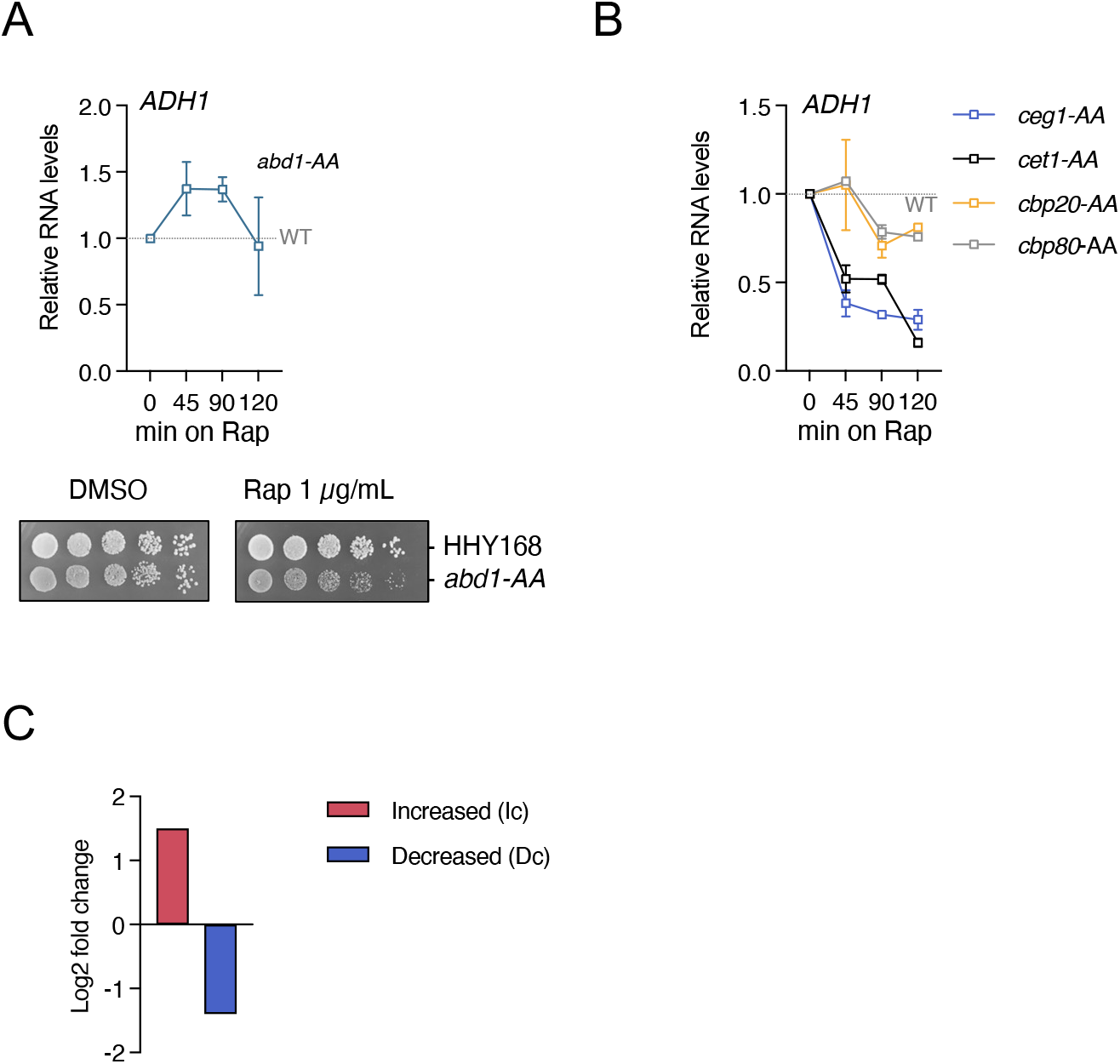
**A.** *ADH1* mRNA levels time during nuclear depletion of Abd1. RT-qPCR RNA levels at 0, 45, 90 and 120 min of rapamycin treatment relative to control (top). The error bars show the standard deviation of three independent experiments. Spot test showing the growth of *abd1-AA* strain on DMSO and Rap (bottom) in comparison with to the parental strain HHY168. Serial dilution of each strain was spotted on YPD media in the presence of DMSO or rapamycin at the final concentration of 1 ug/mL. **B.** *ADH1* mRNA levels during nuclear depletion of Ceg1, Cet1, Cbp20 and Cbp80. RT-qPCR showing the RNA levels measured at 0, 45, 90 and 120 min of rapamycin treatment relative to control (DMSO). The error bars show the standard deviation of three independent experiments. **C.** Bar chart showing the average fold change for decreased (D) and increased (I) mRNA species upon Ceg1 nuclear depletion.

**Figure S2.**
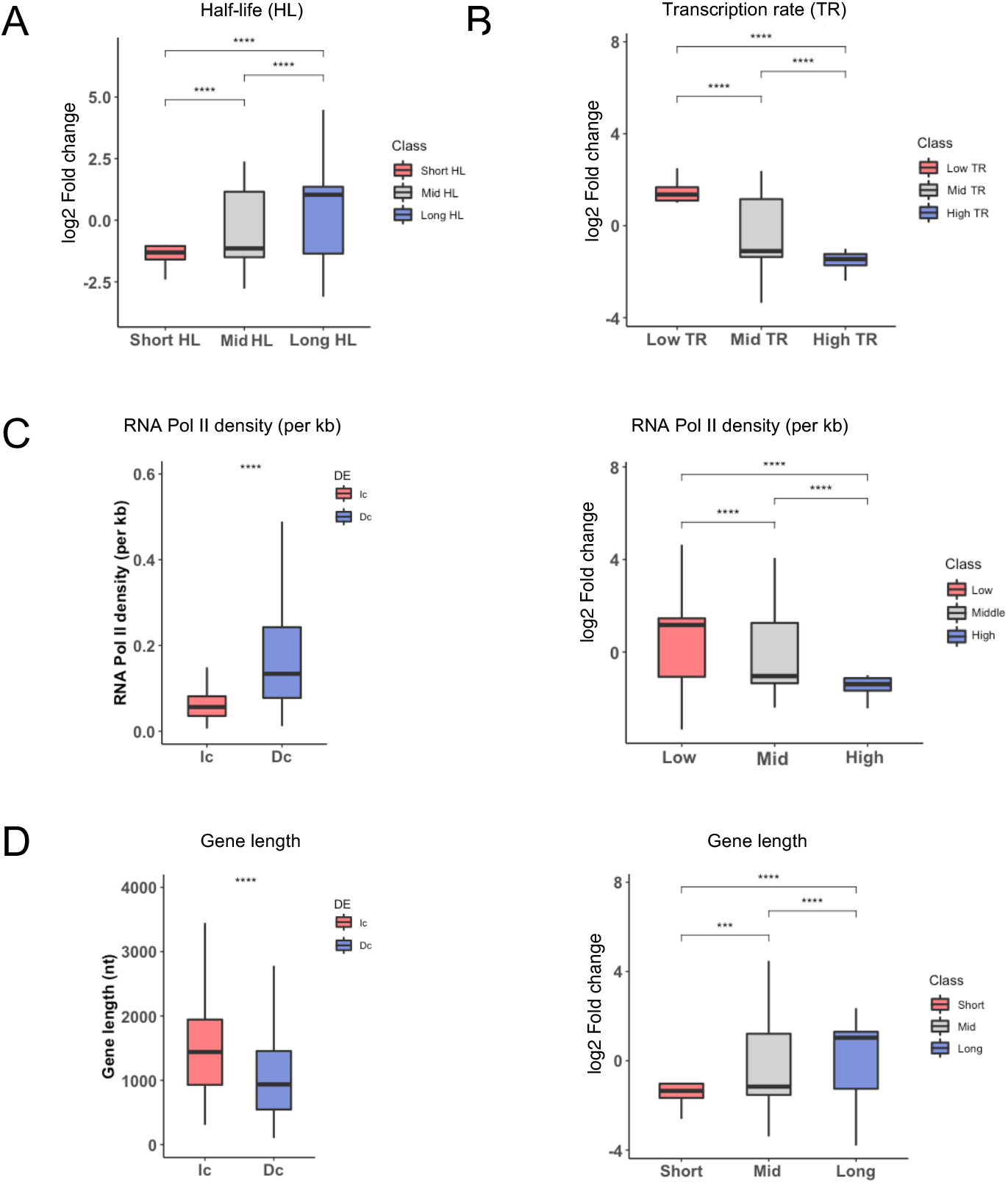
**A-D.** Fold change and fold change sorted by the expression level in *ceg1-AA* (DMSO) for differentially accumulated mRNAs in *ceg1-AA* (Rap) vs *ceg1-AA* (DMSO) plotted accordingly to their half-life (A) and transcription rates (B), Pol II density (C) and gene length (D) sorted by the expression level in *ceg1-AA* (DMSO). ns=P > 0.05; *=P ≤ 0.05; **=P ≤ 0.01; ***= P ≤ 0.001; ****= P ≤ 0.0001.

**Figure S3.**
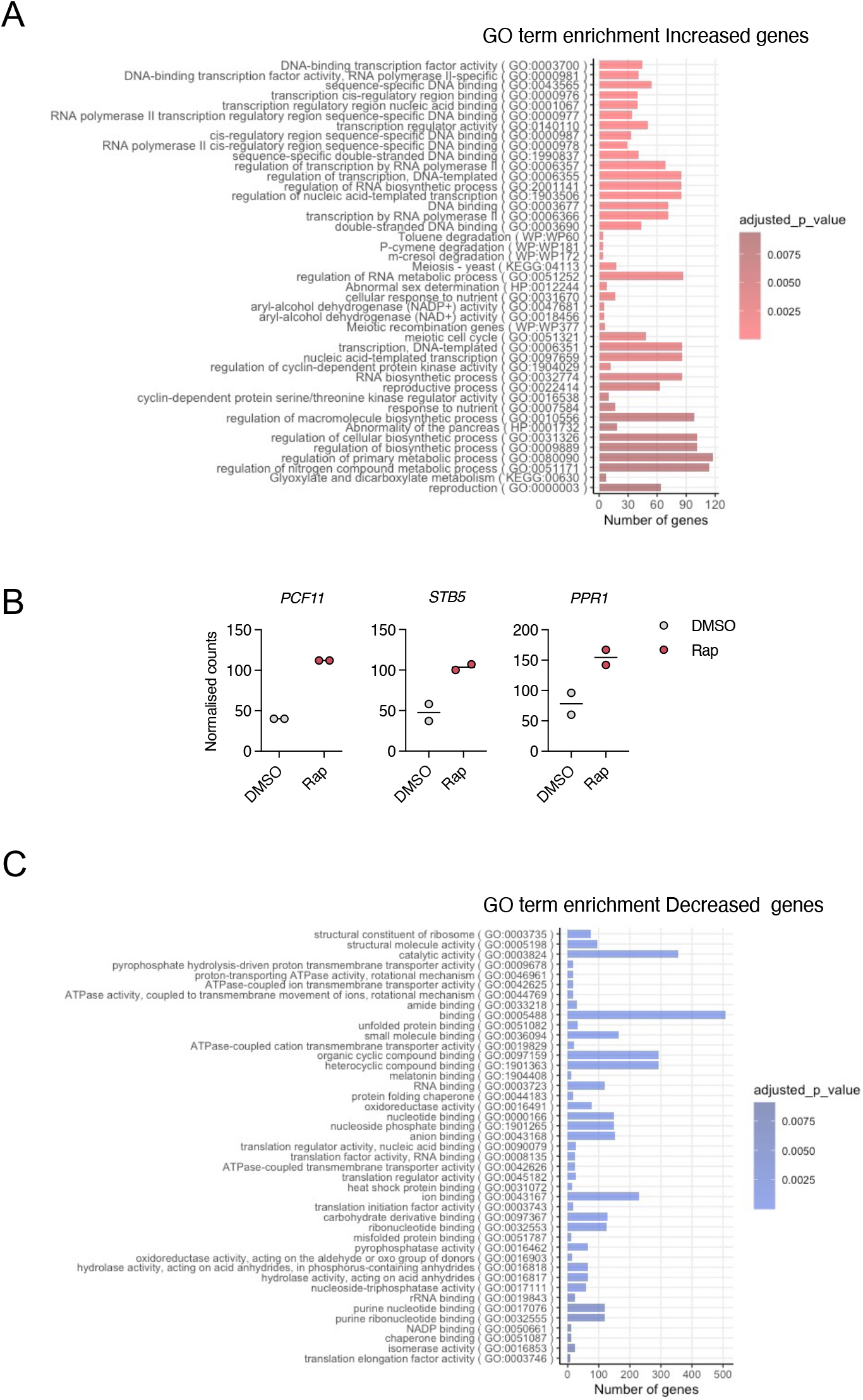
**A.** GO terms associated with increased mRNAs in *ceg1-AA* (Rap). Charts showing the number of the genes belonging to the specific GO group. The classes are listed in a p-value increasing order. **B.** Plots showing the counts per gene for *PCF11, STB5* and *PPR1* obtained from the RNA-seq experiment. Charts showing the number of the genes belonging to the specific GO group. The classes are listed in a p-value increasing order. **C.** GO terms associated with decreased mRNAs in *ceg1-AA* (Rap). Charts showing the number of the genes belonging to the specific GO group. The classes are listed in a p-value increasing order.

**Figure S4.**
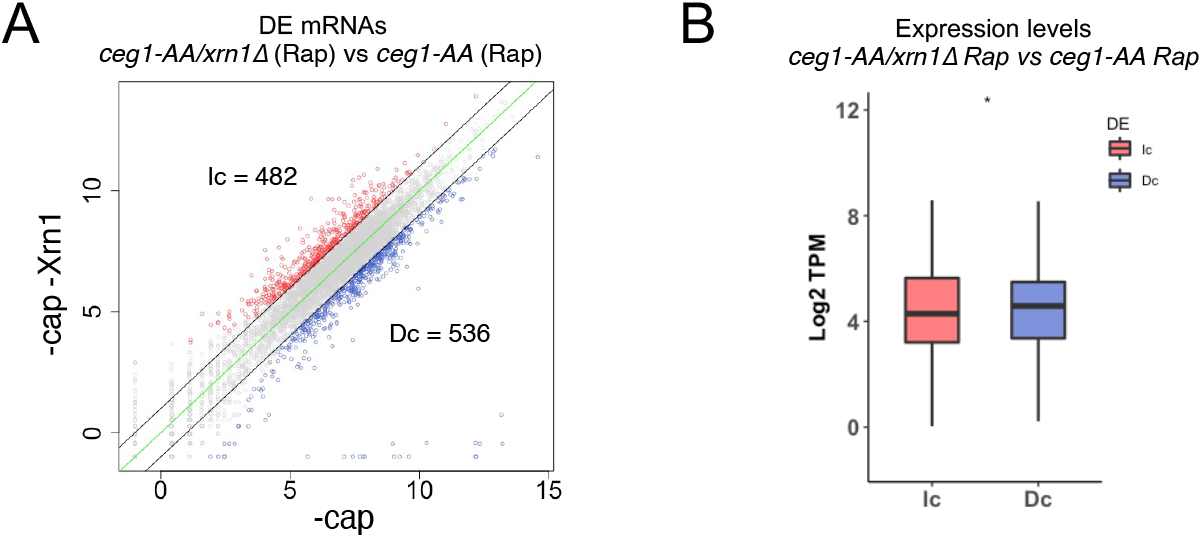
**A.** RNA-seq analysis showing differential accumulation of mRNA species in *ceg1-AA/xrn1D* (Rap) strain after 45 min of rapamycin treatment compared to *ceg1-AA* (Rap). Decreased (D) RNA species showing log2 Fold change <1 are in blue. Increased (I) RNA species with log2 Fold change >1 are labelled in red. The zero-change line is in green. **B.** Differentially expressed mRNAs in *ceg1-AA/xrn1*Δ (Rap) vs *ceg1-AA* (Rap) sorted by their expression levels: Increased (I) or decreased (D). The expression levels were calculated by Transcript Per Million (TPM) value in *ceg1-AA* (DMSO).

**Figure S5.**
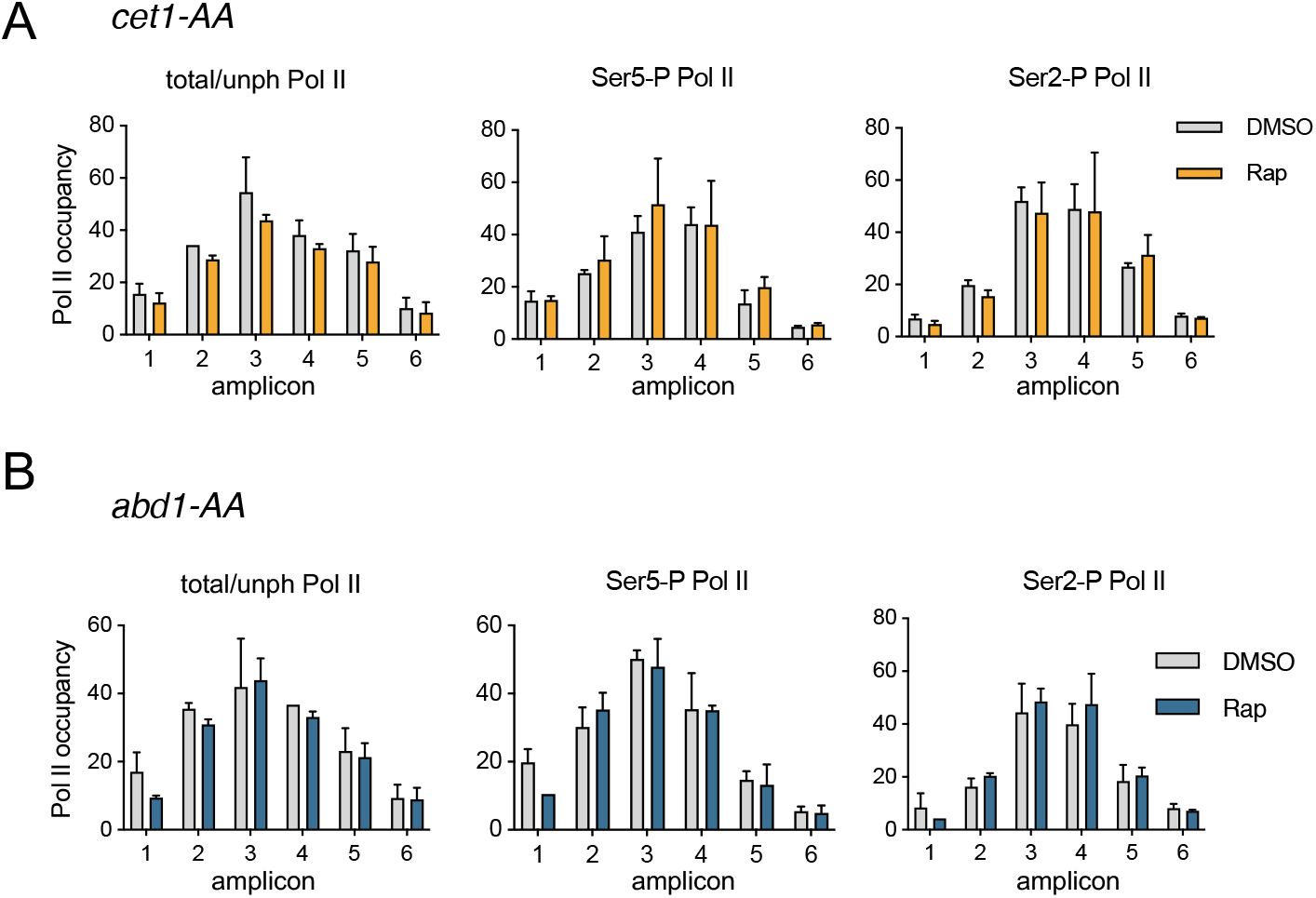
**A-B.** The distribution of total/unphosphorylated (total/unph) Pol II as well as serine 5-phosphorylated (Ser5-P) and serine 2-phosphorylated (Ser2-P) Pol II isoforms over *ADH1* in control (DMSO) and after 45 min of rapamycin (Rap) in (A) *cet1-AA* and *abd1-AA* strain (B). The location of the amplicons used for the ChIP-qPCR is shown above the charts. The error bars show the standard deviation of three independent ChIP-qPCR experiments.

## Notes

### Competing Interest Statement

The authors have declared no competing interest.

### Summary of Updates

A revised version of the manuscript.

